# Ucp2-dependent microglia-neuronal coupling controls ventral hippocampal circuit function and anxiety-like behavior

**DOI:** 10.1101/2020.12.05.413179

**Authors:** Yuki Yasumoto, Milan Stoiljkovic, Jung Dae Kim, Matija Sestan-Pesa, Xiao-Bing Gao, Sabrina Diano, Tamas L. Horvath

## Abstract

Microglia have been implicated in synapse remodeling by phagocytosis of synaptic elements in the adult brain. However, the underlying mechanism of such process is ill-defined. By examining microglia-neuronal interaction in the ventral hippocampus, we found a significant reduction in spine synapse number during the light phase of the light/dark cycle accompanied by increased microglial phagocytosis. This was followed by a transient rise in microglial production of reactive oxygen species (ROS) and uncoupling protein 2 (*Ucp2*) expression, which is a regulator of mitochondrial ROS generation. Conditional ablation of microglial *Ucp2* hindered phasic elimination of spine synapses, increased accumulations of ROS and lysosome-lipid droplet complexes leading to hippocampal circuitry disruption assessed by electrophysiology, and, altered anxiety-like behavior. These observations unmasked a novel and chronotypical interaction between microglia and neurons involved in control of brain functions.

## Introduction

Research on mechanisms behind cognitive and behavioral disorders has historically focused on the structural and functional abnormalities of neurons. Emerging evidence that non-neuronal brain cells, including microglia, are essential in providing metabolic support, modulation of neurotransmitters release and uptake, and regulation of synapse plasticity has uncovered their active involvement in the etiology of these disorders and proposed novel treatment targets (1). Microglia, the resident innate immune cells of the brain, have a pivotal role in the clearance of apoptotic cells and neuronal debris, inflammatory response to pathogens or injury, and in surveillance of local microenvironment. Besides these canonical roles, microglia are important for the remodeling of synaptic connections, an essential process for promoting plasticity and maturation of neural networks during development and adulthood (2). Specifically, microglia through interaction with neurons, dynamically regulate synapse density by modulation of extracellular matrix and phagocytic elimination of synapse (3), (4). This property of microglia also serves to maintain the delicate balance between excitation and inhibition in neuronal assemblies, which is particularly important for the regulation of higher brain functions and effective behavioral control. Indeed, aberrant activity of microglia has been associated with the development of cognitive impairments, depression and anxiety in several neuropsychiatric disorders and with stress (5), (6), (7). However, the underpinning mechanism of microglia-mediated synapse remodeling is still not well understood.

Microglial phagocytosis is followed by a rapid increase in the production of reactive oxygen species (ROS), which facilitate phagocytic contents degradation, but at the same time, may have a detrimental impact on surrounding neurons (8). Phagocytosis is also energetically demanding process that requires high ATP supply. The generations of both ROS and ATP are closely regulated by mitochondria through adaptations in their molecular architecture and signaling. A molecular intermediate involved in these processes is uncoupling protein 2 (*Ucp2*), a member of the mitochondrial transporter superfamily. *Ucp2* is widely expressed in the brain, including in microglia (9), but its functional relevance in the context of microglia-neuronal interaction is ill-defined. To interrogate this, we used recently generated mice with selective deletion of *Ucp2* in microglia (10). Here, we particularly focused on the hippocampus given its profound role in cognition and behavior (11), as well as high immune-vigilant state of microglia (12) and extremely intense spine synapse turnover (13) in this structure of the adult brain.

## Methods

### Experimental animals

Both male and female, adult microglial specific *Ucp2* knock-out mice (*Ucp2*^MGKO^), and their littermate controls (Ucp2+/+-Cx3cr1-cre) were used in the study. *Ucp2*^MGKO^ mice were generated by crossing mice expressing tamoxifen-inducible Cre recombinase (CreERT2) in cells expressing CX3CR1 (Cx3cr1-cre:ERT2) and tdTomato reporter (Ai14; cre recombinase-dependent expression) with mice harboring conditional alleles of *Ucp2* (*Ucp2*fl/fl mice) (10). These mice were injected with Tamoxifen at the age of 5 weeks and all experiments started 4 weeks later to allow the replacement of peripheral monocytes. Before being used, animals were housed in a temperature and humidity-controlled room with a 12:12-h light-dark cycle, with light on at 7 AM: Zeitgeber time (ZT) 0 and off at 7 PM: ZT12. Animals were provided with free access to food and water at all times. During the study, all measures were taken to minimize pain or discomfort of the mice. All procedures were performed according to the protocol reviewed and approved by the Yale University Institutional Animal Care and Use Committee and in compliance with the NIH Guide for the Care and Use of Laboratory Animals (NIH Publications No. 80–23, revised 1996).

### Microglia Isolation

Mice were anesthetized and hippocampi were dissociated from brain and transferred to HBSS buffer. A neural dissociation kit (Miltenyi Biotech, 130-092-628) was used to prepare a single cell suspension, followed by a continuous 30% Percoll gradient at 700g for 15 min. For microglia/macrophage isolation, cells were incubated with biotin conjugated anti-CD11b antibody (eBioscience, 13-0112-82), and CD11b+ cells were isolated using Dynabeads Biotin Binder (Invitrogen, 11047).

### RNA preparation and qPCR

Total RNA was extracted from chopped and homogenized hippocampi, and isolated microglia/macrophage (CD11b+ cells) using RNeasy plus Mini kit (Qiagen, 74134). cDNA synthesis was performed using QuantiTect Reverse Transcription Kit (Qiagen, 205311). qPCR with diluted cDNAs was performed using the LightCycler 480 (Roche) and Taqman Gene Expression Assay primers (Thermo Fisher Scientific) in duplicates. Gene expression of the target genes was normalized to b-actin. Quantification was performed by normalizing Ct (cycle threshold) values to GAPDH and were analyzed by the comparative Ct method with the Roche LightCycler 480 software. The following Primers were used: Ucp2, Mm00627599_m1; b-Actin, Mm02619580.

### Measurement of Mitochondria Oxidation (ROS) in vivo

In vivo mitochondrial ROS production in immunostained microglia was measured by injecting dihydro-ethidium (DHE; Invitrogen by Thermo Fisher Scientific, D11347), as it is specifically oxidized by superoxide to red fluorescent ethidium. A 1mg/ml concentration was injected into retro-orbital sinus of lightly anesthetized mice and mice were transcardially perfused 3 hours later.

### Perfusion and tissue processing

Mice were anesthetized and transcardially perfused with 0.9% saline followed by fixative (4% paraformaldehyde, 15% picric acid, 0.1% glutaraldehyde in 0.1M phosphate buffer (PB; pH 7.4)). Brains were removed and postfixed in the same fixative without glutaraldehyde for 48 hours at 4°C and then stored in 0.1M PB. Coronal 50 µm sections were cut on a PELCO easiSlicer vibratome (TED PELLA, Inc) and followed by immunohistochemistry and electron microscopic analysis.

### Electron microscopy analysis

Free-floating sections (50 µm thick) were incubated with anti-Iba1 antibody diluted 1:3000 in 0.1M PB (Wako Pure Chemical, 019-19741) after 1 hour blocking in 0.1M PB with 5% normal goat serum. After several washes with PB, sections were incubated in the secondary antibody (biotinylated goat anti-rabbit IgG; 1:250 in PB; Vector Laboratories) for 2 hours at room temperature, then rinsed in PB three times 10 min each time, and incubated for 2 hours at room temperature with avidin–biotin–peroxidase (ABC; 1:250 in PB; VECTASTAIN Elite ABC kit, Vector Laboratories, PK6100). The immunoreaction was visualized with 3,3-diaminobenzidine (DAB). Sections were then osmicated (1% osmium tetroxide) for 30 min, dehydrated through increasing ethanol concentrations (using 1% uranyl acetate in the 70% ethanol for 30 min), and flat-embedded in araldite between liquid release-coated slides (Electron Microscopy Sciences, 70880). After embedding in Durcupan (Electron Microscopy Sciences, 14040), ultrathin sections were cut on a Leica Ultra-Microtome, collected on Formvar-coated single-slot grids, and analyzed with a Tecnai 12 Biotwin electron microscope (FEI). To evaluate the ultrastructure within hippocampus striatum radiatum, 10-30 images were taken per mouse. Synapses were counted blindly (expressed as number of synapses per 100 square micrometers), and ImageJ software was used to measure microglial process-synapse contact, phagocytic inclusion coverage, lysosome and LD coverage (expressed as the number per process, percentage of the area to process or cytosol area).

### Immunohistochemistry analysis

Free-floating sections (50 μm-thick) were treated with 0.3% Triton X-100/ PBS. After the sections were blocked in 5% donkey serum, they were incubated with the following primary antibodies for 48 h at 4°C: rabbit anti-Iba1 antibody, 1:2000 (Wako Pure Chemical) and rat anti-CD68, 1:200 (Bio-Rad, MCA1957). The sections were then incubated with the following secondary antibodies for 3 h at room temperature: anti-rabbit IgG-Alexa488 and anti-rat IgG-Alexa488 (1:1000 for both; Invitrogen Co). Sections were mounted on microscope slides and embedded with Vectashield that includes DAPI for nuclear staining (Vector Laboratories, H-1200). Three or four Images of vCA1 per mouse were obtained randomly using confocal microscope (Zeiss LSM880 Airy Scan). A series of 30 μm-thick sections were obtained at 1.0 μm increments along the Z-axis of the tissue section at 20x magnification. The maximum intensity projections (MIP) of the consecutive optical slices were made using Zen software (Zeiss). ImageJ was used for quantification of DHE and CD68 positive area in Iba1 or tomato positive microglia and compared between control and *Ucp2*^MGKO^ mice.

### In vivo electrophysiology

Local field potentials (LFPs) recordings were carried out in mice anesthetized with urethane (1.5 g/kg, i.p.). After achieving a stable plane of anesthesia, which was controlled by the absence of a tail-pinch reflex, mice were placed in a Kopf stereotaxic frame (Tujunga, CA) on a temperature-regulated heating pad (Physitemp Instruments Inc., Clifton, NJ) set to maintain body temperature at 37-38 °C, and unilateral craniotomies were performed above the hippocampus. After surgery, a bipolar concentric stainless steel recording electrodes (NE-100X, Rhodes Medical Instruments, Woodland Hills, CA) were inserted into the dorsal and ventral aspect of hippocampal CA1 region, and animals were allowed to stabilize for 30 min before the beginning of LFPs recordings. Coordinates for positioning electrodes were taken from mouse stereotaxic brain atlas (14) and referenced relative to bregma and brain surface (dorsal CA1 (dCA1): 2.0 mm posterior, 1.5 mm lateral and 1.5 mm dorsoventral, and ventral CA1 (vCa1): 3.4 mm posterior, 3.5 mm lateral, and 3.5 mm dorsoventral). Throughout the duration of the experiment, mice were kept in the stereotaxic frame, spontaneous LFPs were continuously monitored, and the level of anesthesia regularly checked. After all recordings, animals were deeply anesthetized, transcardially perfused, and their brains removed for histological analysis.

In each experiment, LFPs were amplified and filtered between 1 and 300 Hz using Grass P55 AC differential amplifier (Grass Technologies, West Warwick, RI) with an additional notch filter at 60 Hz. The signal was simultaneously sampled at a rate of 1 kHz and stored on a computer via a CED Micro1401-3 interface and Spike2 software (Cambridge Electronic Design, Cambridge, UK).

For quantitative offline analyses of LFPs power, signal was subjected to Fast Fourier transform (FFT) at a spectral resolution of 0.24 Hz. Power analyses during spontaneous hippocampal activity were performed using 10-minute long epochs by summing FFTs in theta (4-9 Hz), and gamma (30-90 Hz) frequency range. For gamma power computation, LFPs signal was first band-pass filtered between 30 Hz and 90 Hz. In subset of mice effect of NMDA receptors blockade on gamma oscillation in vCA1 was tested. After recording of stable baseline of at least 30 min, mice were subcutaneously injected with 0.5 mg/kg MK-801 (Tocris Bioscience, Bristol, UK) and LFPs followed for an additional 60 min. Drug induced changes in gamma power were calculated as the percent change from baseline power in every animal over 10 min post-treatment interval, starting 30 min following injection.

### In vitro electrophysiology

Coronal hippocampal slices of 300 µm thickness, were routinely prepared from 3-week old mice. Briefly, wild type and mutated mice were deeply anesthetized with isoflurane and decapitated, their brains rapidly removed, trimmed to a tissue block containing only the cortex and hippocampus. Tissue was sectioned using vibratome in an oxygenated (5% CO_2_ +95% O_2_) cutting solution at 4°C containing (in mM): sucrose 220, KCl 2.5, CaCl_2_ 1, MgCl_2_ 6, NaH_2_ PO_4_ 1.25, NaHCO_3_ 26, and glucose 10, pH 7.3 with NaOH. After preparation, slices trimmed to contain only the hippocampus were maintained in a storage chamber with oxygenated artificial cerebrospinal fluid (ACSF) containing (in mM): NaCl 124, KCl 3, CaCl_2_ 2, MgCl_2_ 2, NaH_2_ PO_4_ 1.23, NaHCO_3_ 26, glucose 10, pH 7.4 with NaOH. Hippocampal slices were transferred to a recording chamber constantly perfused with ACSF at 33 °C with a rate of 2 ml/min after at least an hour of recovery. Whole-cell voltage clamp (at −60 mV or at 0 mV) was performed as described previously (15) to observe miniature excitatory and inhibitory postsynaptic currents (mEPSC and mIPSC) using Multiclamp 700A amplifier (Axon Instruments, CA). The patch pipettes with a tip resistance of 4-6 MΩ were made of borosilicate glass (World Precision Instruments) with a Sutter pipette puller (P-97) and filled with a pipette solution containing (in mM): K-gluconate 135, MgCl_2_ 2, HEPES 10, EGTA 1.1, Mg-ATP 2, Na_2_ -phosphocreatine 10, and Na_2_ -GTP 0.3, pH 7.3 with KOH. After the giga-ohm (GΩ) seal and whole-cell access were achieved, the series resistance (10-20 MΩ) were partially compensated by the amplifier. All data were sampled at 10 kHz and filtered at 3 kHz with an Apple Macintosh computer using AxoGraph X (AxoGraph, Inc). Data were analyzed using an event-detection package in AxoGraph X and plotted with Igor Pro software (WaveMetrics, Lake Oswego, OR).

### Behavioral tests

All behavior tests were performed in a dimly lit room during dark period and tracked with Any-Maze software (Stoelting Co., Wood Dale, IL). For acclimation, mice were transferred to examination room at least 1 h prior to the tests.

### Elevated Zero Maze

Anxiety-like behavior was assessed using an elevated zero maze consisted of a ring-shaped platform, 60 cm in diameter, 4.3 cm wide, and elevated 52 cm above the floor. The maze was divided into four equal quadrants, two opposing arms were enclosed by 15.5cm high gray walls whereas two were open. Each mouse was placed in center of an open arm and allowed to explore the platform for 5 minutes. Mouse behavior was recorded with an overhead video camera connected to tracer software (ANY-maze; Stoelting Company). The time spent and distance traveled in the open arm were measured by the ANY-maze Software.

### Novel Object Recognition Test

After 24 hours habituation to an open field (37 cm×37 cm×30 cm), each mouse was presented with two exact same objects (Falcon tissue culture flasks filled with corn-based animal bedding (10 cm high, 2.5 cm deep and 5.5 cm wide, transparent plastic with a blue bottle cap)) in the field and allowed to explore for 10 minutes. After a short intersession interval (2 hours), each mouse was placed back to the same box, where one of the two familiar objects was switched to a new one (towers of Lego bricks (10 cm high and 5.8 cm wide, built in blue, yellow, red, purple and green bricks)). Mouse behavior was recorded with an overhead video camera connected to ANY-maze. The amount of time taken to explore the new object was measured by the ANY-maze Software and the novel preference and the discrimination index percentage were calculated by the following formula. Novel preference (%)=Time exploring novel object/(Time exploring novel object+Time exploring familiar object)×100. Discrimination index (%)=(Time exploring novel object–Time exploring familiar object)/(Time exploring novel object+Time exploring familiar object)×100.

### Marble Burying Test

The cages were filled with 5 cm of corn-based animal bedding. Twenty four glass marbles were evenly placed on the surface and leave each mouse for 30 minutes. The number of marbles buried more than 2/3 their depth with bedding was counted.

### Y Maze

A Y maze apparatus has three arms (30 cm×30 cm×30 cm), 8 cm lane width and 15 am wall height at 120° angle from each other. Each mouse was placed at the end of one arm facing the center and allowed to freely explore the three arms for 10 minutes. Mouse behavior was recorded with an overhead video camera connected to ANY-maze. The number of triads was defined as consecutive entries into each of the three arms without repetition and scored as alteration. The percentage of alteration was calculated by the following formula. Alteration (%)=The number of alterations/(Total arm entries-2)×100

### Statistical analysis

All data were initially determined to be suitable for parametric analysis according to normality and homoscedasticity. Statistical difference was assessed using one-way ANOVA with Tukey’s post hoc test and two-tailed Student’s t-test (GraphPad Prism Software, Inc, La Jolla, CA). Differences were considered significant when p < 0.05.

## Results

### Spine synapse number and morphology are regulated in a chronotypical manner in the ventral hippocampal CA1 region

First we found that microglia-mediated spine synapse elimination in the hippocampus exhibits a chronotypical pattern with a higher propensity for synaptic phagocytosis during the light phase. Electron microscopic (EM) analyses revealed a significant decrease in the number of spine synapses in the stratum radiatum of the ventral hippocampal CA1 region (vCA1) of male mice at ZT6 (the middle of the light phase) comparing to ZT0, ZT12, and ZT18 time points (F_(3,116)_ =32.10, P<0.0001). In contrast, post-synaptic morphological features i.e. post-synaptic density (PSD) length (F_(3,536)_ =37.23, P<0.0001), post-synaptic head area (F_(3,536)_ =6.33, P<0.0001), diameter (F_(3,536)_ =5.80, P=0.0007) and length (F_(3,536)_ =7.07, P<0.0001) of these synapses were all markedly larger only at ZT18 (the middle of the dark phase), (Fig. 1A-D, Table 1). The same pattern was observed in female mice (F_(3,116)_ =27.30, P<0.0001) (Supplemental Fig. 1A).

**Table 1.**
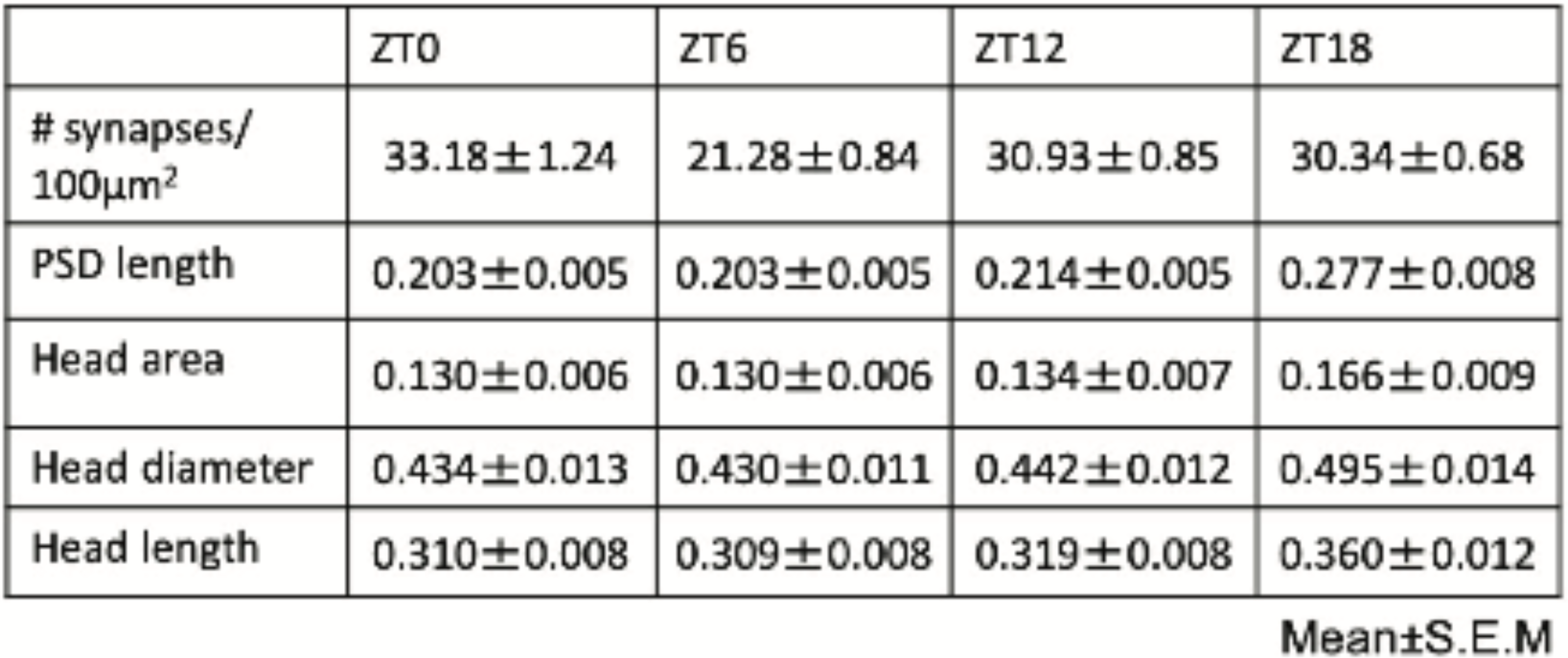
Spine synapse number counts and postsynaptic morphological characteristics of vCA1 region in control male mice at different zeitgeber time periods (ZT0-ZT18). Data are expressed as mean ± s.e.m.

|  | ZT0 | ZT6 | ZT12 | ZT18 |
| --- | --- | --- | --- | --- |
| # synapses/<br>100$\mu\text{m}^2$ | 33.18±1.24 | 21.28±0.84 | 30.93±0.85 | 30.34±0.68 |
| PSD length | 0.203±0.005 | 0.203±0.005 | 0.214±0.005 | 0.277±0.008 |
| Head area | 0.130±0.006 | 0.130±0.006 | 0.134±0.007 | 0.166±0.009 |
| Head diameter | 0.434±0.013 | 0.430±0.011 | 0.442±0.012 | 0.495±0.014 |
| Head length | 0.310±0.008 | 0.309±0.008 | 0.319±0.008 | 0.360±0.012 |
| Mean±S.E.M | | | | |

**Fig. 1.**
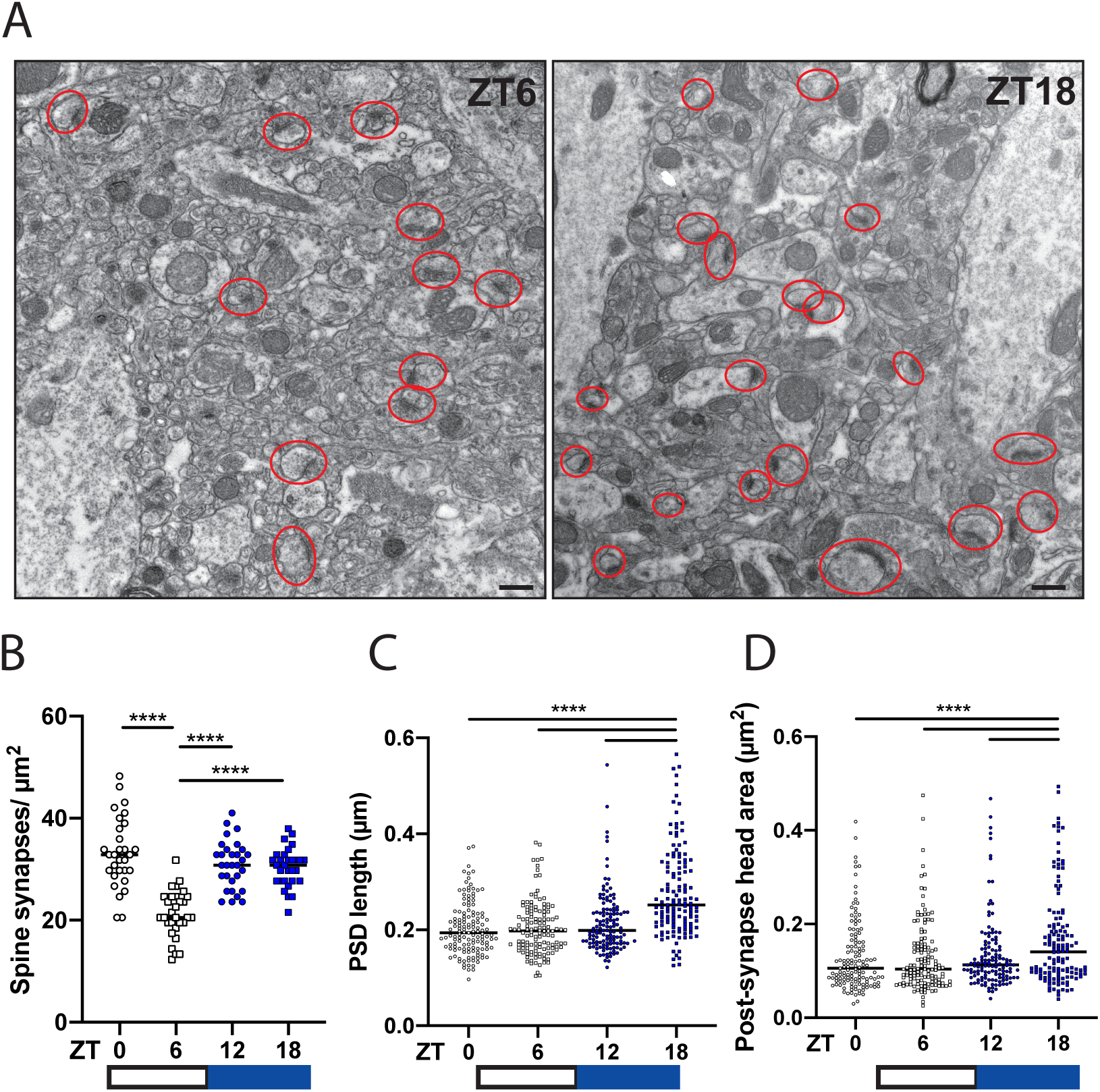
Chronotypical pattern of spine synapse number and morphological alterations in vCA1 region. **A-D**, Representative electron microscopic images of spine synapses (encircled in red) in stratum radiatum layer of vCA1 of male mice at zeitgeber times ZT6 (left) and ZT18 (right) (**A**). Dynamic changes of synapse number (**B**), postsynaptic density (PSD) (**C**), and postsynaptic head area (**D**) at ZT0, ZT6, ZT12 and ZT18. Data are presented as scatter plot graphs of replicates (n=3 mice/group, 10 images/mouse (**B-D**)) with medians indicated by horizontal lines (**B-D)**. ****p<0.0001, by one-way ANOVA with Tukey’s post hoc test (**B-D)**. Scale bars, 500 nm (**A**).

### Chronotypically regulated microglial synaptic pruning induced ROS production and UCP2 expression

Next to assess microglial engulfment activity, we quantified microglia-synapse contacts per process and relative phagocytic inclusion coverage by EM. We found a significantly increased contact number (t=5.34, df=60, P<0.0001) and inclusion area (t=5.29, df=60, P<0.0001) during the light phase (Fig. 2A-D). Further, we detected rise in microglial ROS production peaking at ZT12 (the end of the light phase) (F_(3,427)_ =31.67, P<0.0001) (Fig. 2E, F), followed by subsequent upregulation of *Ucp2* mRNA expression in the bulk hippocampus (Fig. 2G), and in isolated hippocampal microglia (Fig. 2H) during the dark phase (t=3.66, df=8, P=0.006; t=3.22, df=7, P=0.015, respectively). Together, these results indicate that microglia is actively involved in synapse remodeling in mice vCA1 during their daylight dormancy and this process is accompanied by increased ROS and *Ucp2* levels.

**Fig. 2.**
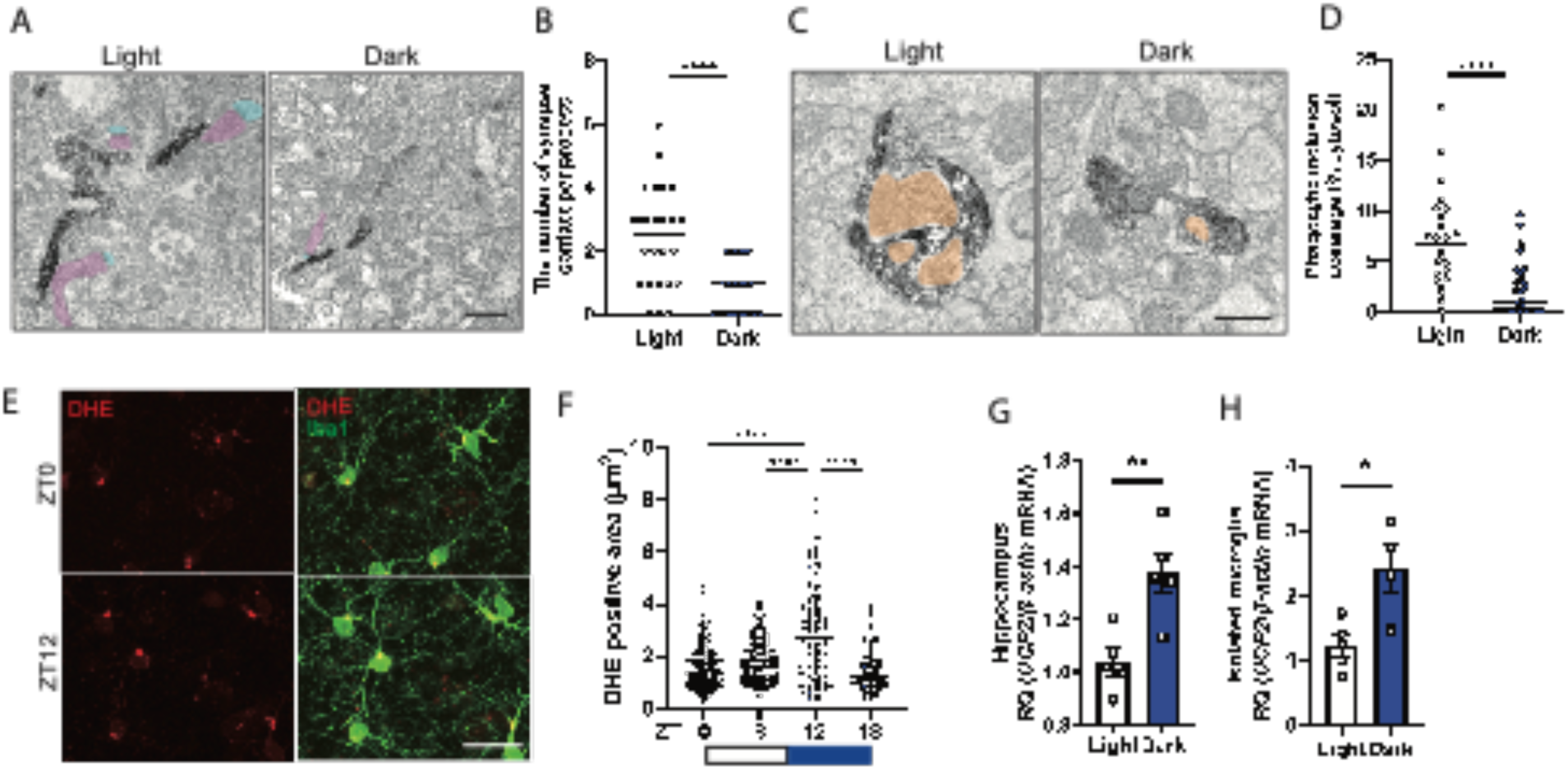
Chronotypical pattern of microglial phagocytic activity in vCA1 region. **A-D** Representative electron microscopic images and quantification of microglia-synapse contacts per process (**A, B**), and phagocytic inclusion coverage (**C, D**) at light (left) and dark (right) phase. Immunostained Iba1 sections showing presynaptic (pink), postsynaptic (blue) compartments and phagocytic inclusions (orange). **E, F**, Representative confocal images of DHE+ (red) and Iba1+ (green) at ZT0 and ZT12 (**E**), and quantification of DHE+ area in Iba1 immunoreactive microglia (**F**). **G, H**, Quantitative PCR analyses of relative *Ucp2* mRNA expression in the bulk of hippocampus (**G**) and isolated hippocampal microglia (**H**) at light and dark phase. Data are presented as scatter plot graphs of replicates (n=3 mice/group, 3-4 images/mouse (**F**) 8-12 images/mouse (**B**,**D**)) with medians indicated by horizontal lines (**B, D, F**), and histograms with individual values (n=5 mice/group) and s.e.m. (**G, H**). *p<0.05; **p<0.01;****p<0.0001, by one-way ANOVA with Tukey’s post hoc test (**F**), or two-tailed Student’s t-test (**B, D, G, H**). Scale bars, 500 nm (**A, C**), 20 μm (**E**).

### Loss of Ucp2 in microglia leads to increased ROS production and aberrant phagocytic phenotype

To further determine *Ucp2* functional role in the regulation of microglial phagocytosis-mediated synapse remodeling and plasticity in the hippocampus, we used *Ucp2*^MGKO^ mice with microglial specific ablation of *Ucp2*. To detect the ROS level in these mice, dihydroethidium (DHE) was injected intravenously and quantified DHE positive area. In addition, we used CD68, a marker of lysosome, to quantify lysosome accumulation. We found significantly increased ROS level (t=5.35, df=142, P<0.0001) and accumulation of lysosomes (t=4.49, df=206, P<0.0001) in microglia of hippocampal vCA1 compared to their controls at ZT12 (Fig. 3A, B). Moreover, mutant mice had larger lysosomal vacuoles and a higher number of lipid droplets covering a sizeable portion of the microglial cell body and processes (t=2.02, df=53, P=0.049) in EM analyses (Fig. 3D, E). Also, these mice showed significantly fewer synapse contacts (t=3.06, df=43, P=0.004), and phagocytic inclusions (t=4.19, df=52; P<0.0001) in microglial processes comparing to controls (Fig. 3F, G).

**Fig. 3.**
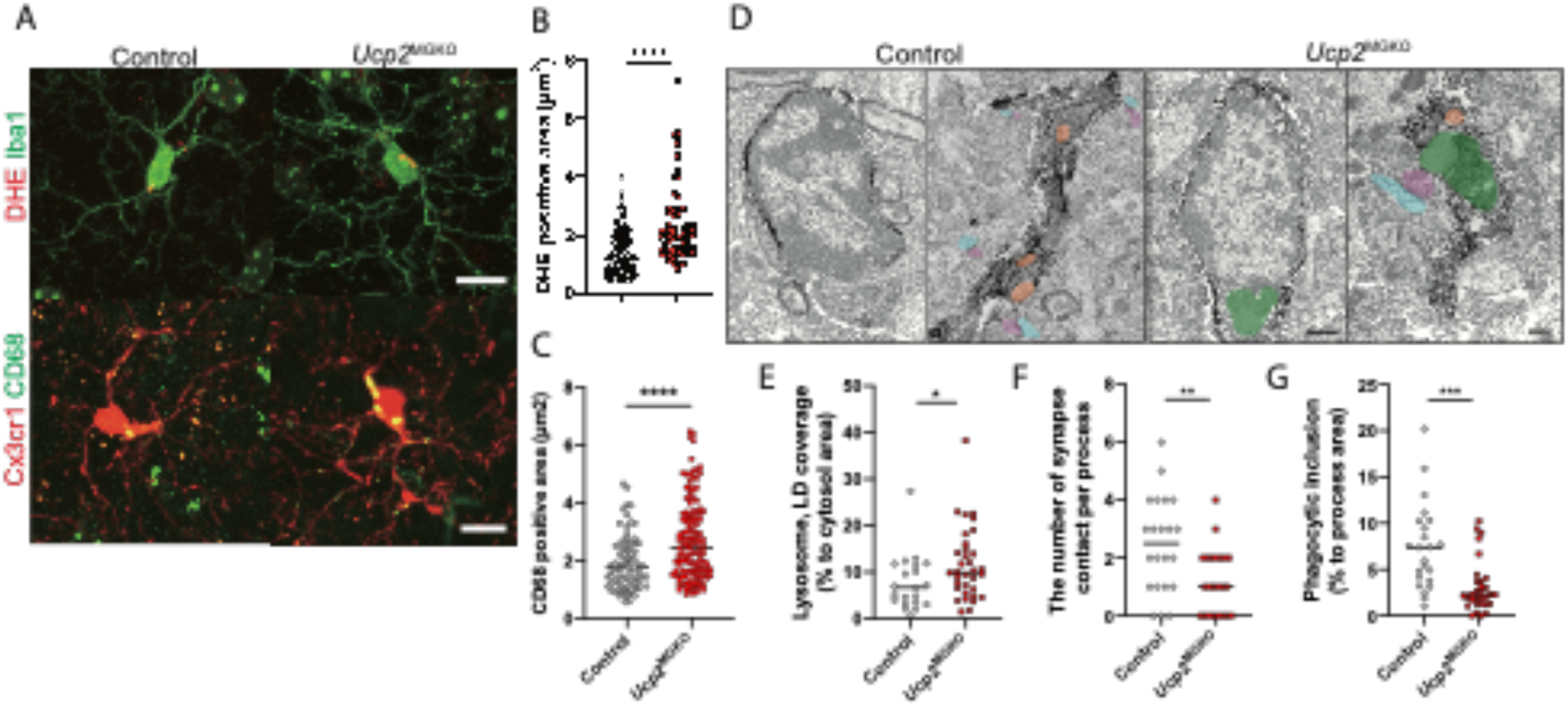
Alteration of microglial ROS production, phagocytosis and synapse number in vCA1 region by microglia selective *Ucp2* deletion. **A-C**, Representative confocal images showing DHE probes (red) and Iba1 staining (green) of control (left) and *Ucp2*^*MGKO*^ (right) male mice (**A**, upper panels), and tomato-expressing microglia (red) and CD68 staining (green) (**A**, lower panels). Quantification of DHE+ area in Iba1 immunoreactive microglia (**B**) and CD68+ area (**C**). **D-G**, Representative electron microscopic images of Iba1-immunostained microglial cell body (upper panels) and process (lower panels) of control (left) and *Ucp2*^*MGKO*^ (right) male mice (**D**). Quantification of lysosomal and lipid droplet coverage (**E**), number of synapse contact per microglial process (**F**) and phagocytic inclusion area (**G**). Data are presented as scatter plot graphs of replicates (n=3 mice/group, 5-10 images per animal) with medians indicated by horizontal lines (**B, C, E-G, B**). *p<0.05; **p<0.01; ***p<0.001; ****p<0.0001, by two-tailed Student’s t-test (**B**,**C, E-G**). Scale bars, 10 μm (**A**), 500 nm (**D**).

### Microglial specific UCP2 deletion induced increase in synapse number in the ventral hippocampal CA1 region

As a consequence, *Ucp2*^MGKO^ mice had a remarkably higher number of spine synapses in the stratum radiatum of vCA1 measured over ZT0-ZT12 periods (F_(7,232)_ =15.28, P<0.0001) (Fig. 4A, B). However, temporal dynamics of synapse elimination therein was still preserved (F_(3,116)_ =41.46, P<0.0001) (Fig. 4B). By examining synapse morphology in *Ucp2*^MGKO^ mice we detected shorter PSD length and smaller post-synapse area than in control littermates (data not shown). Importantly, these synapse alterations were not observed in the dorsal CA1 region (dCA1) of *Ucp2*^MGKO^ mice (Supplemental Fig. 2A), suggesting that *Ucp2* may play a role in the region-specific function of microglia within the hippocampus and brain.

**Fig. 4.**
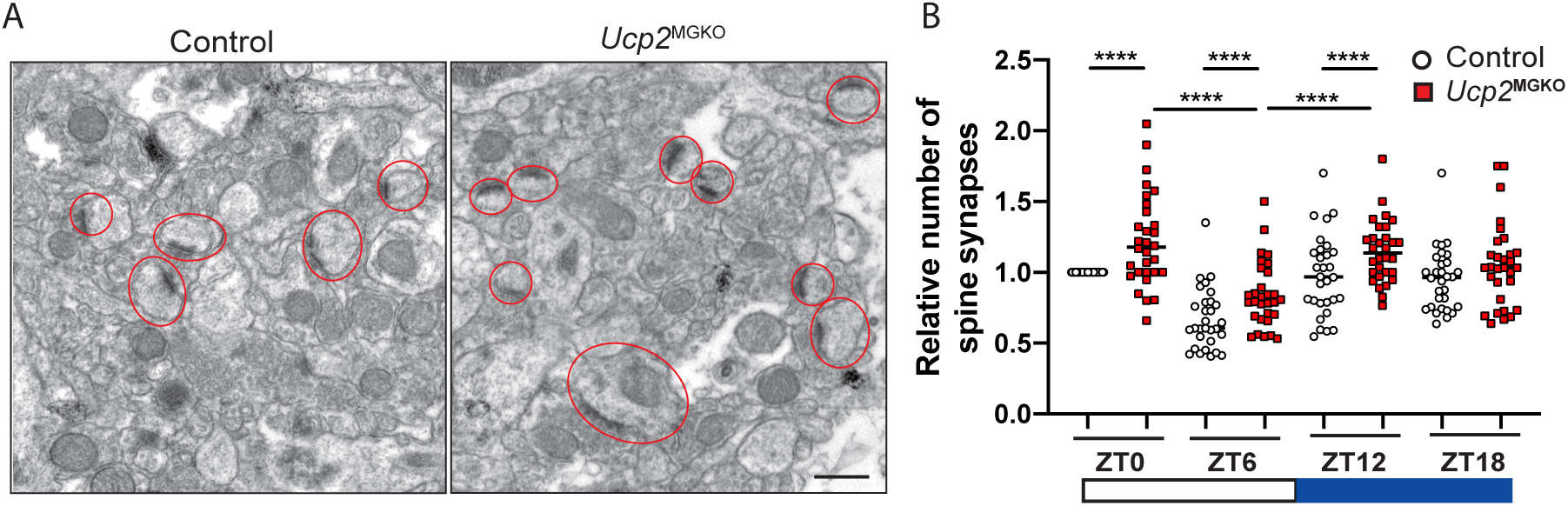
Alteration of synapse number in vCA1 region by microglia selective *Ucp2* deletion. **A**,**B**, Representative electron microscopic images of spine synapses (encircled in red) in stratum radiatum layer of vCA1 of control (left) and *Ucp2*^*MGKO*^ (right) male mice (**A**) with relative spine synapse counts at ZT0 (**B**). Data are presented as scatter plot graphs of replicates (n=3 mice/group, 10 images per animal) with medians indicated by horizontal lines (**B**). ****p<0.0001, by one-way ANOVA with Tukey’s post hoc test (**B**). Scale bars, 500 nm (**A**).

### Microglial specific UCP2 deletion altered neurophysiological functions and anxiety-like behavior

To interrogate the effect of *Ucp2*-dependent alterations of microglial synapse remodeling on hippocampal circuit integrity we performed both *in vivo* and *in vitro* electrophysiological recordings from male and female *Ucp2*^*MGKO*^ and control mice. Recordings of LFPs from the vCA1 in anesthetized *Ucp2*^MGKO^ male mice revealed a significant reduction in gamma oscillation power (t=2.61, df=12, P=0.029) and suggestive decline in theta oscillation power (t=1.81, df=10, P=0.099) (Fig. 5A-C). Interestingly, systemic administration of MK-801, a noncompetitive antagonist of glutamatergic NMDA receptor, was able to rescue this altered gamma oscillation given the comparable increase in vCA1 gamma power in both *Ucp2*^MGKO^ and control mice (t=1.04, df=4, P=0.359) (Fig. 5D). In contrast, changes in these oscillatory bands were not found in dCA1 of male *Ucp2*^*MGKO*^ mice comparing to their control counterparts (Supplemental Fig. 2B-D). Additionally, female *Ucp2*^MGKO^ mice also showed similar results as controls for hippocampal gamma and theta power in both vCA1 and dCA1 recordings (Supplemental Fig. 1B-E). In further characterization of neuronal alteration observed in vCA1, we performed whole-cell voltage clamp recordings in hippocampal slices and compared miniature excitatory/inhibitory postsynaptic currents (mEPSC/mIPSC) in male *Ucp2*^MGKO^ and control mice. We found significantly higher frequency of mIPSC in *Ucp2*^MGKO^ (t=2.24, df=25, P=0.035) (Fig. 5E, F), which confirmed functional impairment of neuronal activity in vCA1 in these mice. Finally, behavioral testing of *Ucp2*^MGKO^ mice revealed increased anxiety level. In the elevated zero maze test, male mutant mice spent significantly less time (t=2.50, df=17, P=0.023) (Fig. 5G) and had shorter distance traveled (t=2.11, df=17, P=0.050) (Fig. 5H) in the stressful open arms as compared to their controls. In contrast, this anxiety-like phenotype was not observed in female mice (Fig. 5G, H). Moreover, comparing performance in novel object recognition, marble burying, and Y-maze tasks between *Ucp2*^MGKO^ and control animals of either gender did not reveal any obvious alteration (Supplemental Fig. 3A-D).

**Fig. 5.**
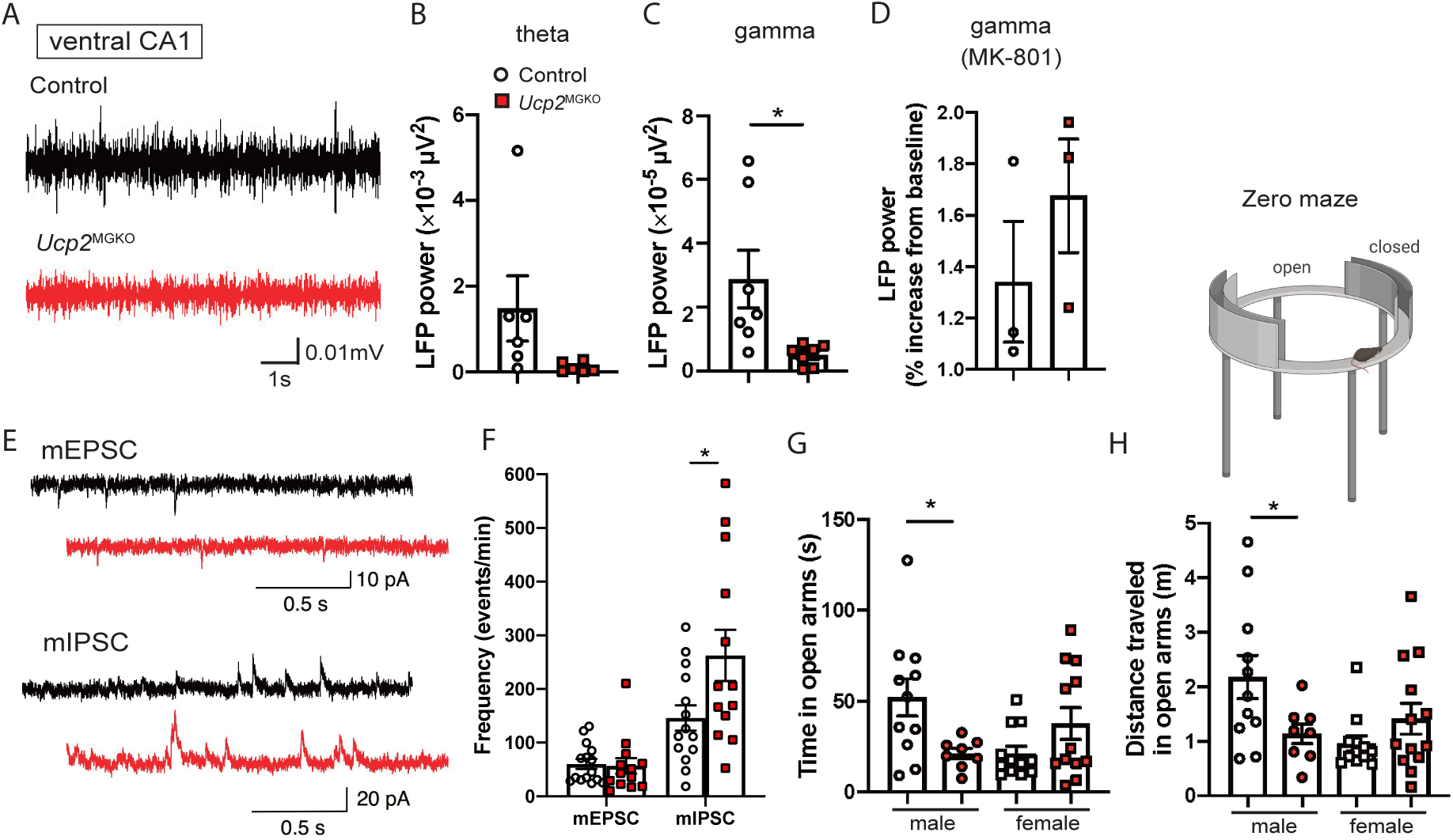
Neurophysiological dysfunctions and behavioral alterations in *Ucp2*^*MGKO*^ male mice. **A-D**, Typical traces of spontaneous local field potentials (LFPs) in gamma frequency range from control (black) and *Ucp2*^*MGKO*^ (red) male mice recorded *in vivo* from vCA1 region. Both signals are digitally band-pass filtered between 30 and 90 Hz (**A**). Power of theta (4-9 Hz) (**B**), and gamma (30-90 Hz) (**C**) band oscillations. Relative change in gamma power following MK-801 (0.5 mg/kg) subcutaneous injection (**D**). **E-F**, Representative traces of miniature excitatory postsynaptic current (mEPSC, upper panel) and miniature inhibitory postsynaptic current (mIPSC, lower panel) from control (black) and *Ucp2*^*MGKO*^ (red) mice acquired from whole-cell recordings in vCA1 (**E**), and corresponding analysis of spontaneous mEPSC and mIPSC frequency of occurrence (**F**). **G-H**, Comparisons of time spent (**G**) and distance traveled (**H**) in open arms of elevated zero maze test between male and female control and *Ucp2*^*MGKO*^ mice. Data are presented as histograms with individual values and s.e.m. (n=5 mice/group for **B**; n=7 mice/group for **C**; n=3 mice/group for **D** and **F**; n=8-12 mice/group for **G** and **H**). *p<0.05, by two-tailed Student’s t-test.

## Discussion

In the healthy adult brain, microglia-neuronal interaction is believed to have a prominent role in securing permanence of neuronal activity in the optimal range, thereby providing homeostatic control of network function and behavior. Here, we show that this interaction in the mouse ventral hippocampus operates in the chronotypical pattern with microglia active phagocytic pruning of synapses during the light phase. We provide evidence that this process critically depends on mitochondrial protein *Ucp2*, since ablation of *Ucp2* in microglia hindered synapse elimination triggering changes in hippocampal circuit function and inducing anxiety-like behavior.

Alternating environmental states of day and night are considered among the strongest salient stimuli capable to entrain changes in neuronal structure and function that confer synaptic plasticity (16). In the hippocampus, the magnitude of plasticity in the CA1 region is regulated in a time-dependent manner, and this may help prevent saturation of long-term potentiation/depression, and the hyperstimulation of local networks, thereby limiting their negative impact on cognition and neuronal health (17). These important aspects of homeostasis are partly gated by circadian rhythm-driven expression of hippocampal clock gene (16), but other mechanisms including microglial activation as we observed may also have substantial involvement. In support of the latter, we found that synapse phagocytosis in vCA1 is followed by a rise in microglial ROS production which is known to enable phagocytic clearance. However, at the same time ROS excessive formation can cause cell damage and reduce engulfment capacity. Previous evidence showed that *Ucp2* can toggle ROS level protecting cells from excessive ROS formation (8), (18). Consistently, we detected significant increase of *Ucp2* both in the hippocampus and isolated hippocampal microglia during the dark phase after the peak of ROS production.

In peripheral macrophages, *Ucp2* is a major molecular determinant that mediates mitochondrial function during phagocytosis (18). Recently we provided evidence that *Ucp2*-dependent mitochondrial changes contribute to microglial activation and neuroinflammation induced by high-fat diet (10). In current study, we show that microglial *Ucp2* plays an important role in lysosomal-autophagy pathway since analysis of *Ucp2*^MGKO^ mice lacking microglial Ucp2 revealed both significant increase in ROS, and defective lysosomal degradation of lipid droplets. It has been suggested that accumulated lipid droplets in microglia contribute to impairment of cellular energy metabolism given the shortage in free fatty acids for mitochondrial β-oxidation and generation of ATP provided by their degradation (19). Moreover, lipid droplets themselves may induce inflammation and sequestration of lysosomes compromising their role in phagocytosis as shown for the hippocampal microglia after lipopolysaccharide treatment or during aging (19). Both these could propel into further microglial dysfunctions. Accordingly, *Ucp2*-driven substantial transformation of microglia in *Ucp2*^MGKO^ mice resulted in suppression of spine synapse remodeling that yield to alteration in hippocampal circuit function and related behavior.

In vivo recordings of LFPs revealed reduction in gamma oscillation power in vCA1 of *Ucp2*^MGKO^ male mice, without marked changes in activity in dCA1 or in hippocampus of female knock-out mice. It is well-established that network oscillations depend on the integrity of the synaptic contacts and precise temporal synchronization of neural activity in local circuits (20). Furthermore, high-frequency oscillation phenomena, such as gamma (30-90 Hz), greatly rely on fine-tuning of excitation and inhibition between pyramidal cells and interneurons (21). Given that spine synapses closely correspond to the excitatory synapses (22), their increase in number as found in *Ucp2*^MGKO^ mice, can induce an excitatory gain in the local vCA1 circuitry which offset the excitatory/inhibitory balance. This results in asynchronous firing of pyramidal neurons reflected as decrease in oscillation power. In the hippocampal network during the gamma cycle, phasic glutamate excitatory inputs from pyramidal neurons, via feedback loop drive parvalbumin-positive (PV+) interneurons to fire with a delay due to the monosynaptic nature of recurrent connections (21). Since gamma oscillation critically depends on synchronous activation of inhibitory fast-firing PV+ interneurons, increasing excitation in synapses onto these cells, which can be long-lasting in case of glutamate NMDA mediated-currents, prompts PV+ interneurons to generate action potentials that are not locked to the excitatory discharge from the pyramidal cells inducing firing desynchronization in network. In agreement, computational model network of reciprocally connected pyramidal cells and interneurons demonstrated that increasing NMDA tone decrease gamma power, suggesting importance of NMDA receptors in driving inhibitory interneuron activity (23). Indeed, pharmacological blocking of NMDA receptors increased gamma power in vCA1 of *Ucp2*^MGKO^ mice to the level of control mice. This result aligns with findings of enhanced spontaneous hippocampal gamma oscillation in mice with selective ablation of NMDA receptors located onto PV+ interneurons (24). Additionally, our analysis of higher mIPSC in vCA1 neurons of *Ucp2*^MGKO^ mice further corroborate alteration in inhibition.

Recent study showed that microglia can suppress neuronal activity in response to extracellular ATP released upon neuronal activation. This negative feedback mechanism is region-specific and likely functions similarly to inhibitory neurons to constrain excessive excitation in the brain (25). Our findings of microglial impact on shaping inhibitory output in neural circuitry by spine synapse regulation extend this notion indicating that these two microglia-driven mechanisms might cooperate synergistically to maintain excitatory/inhibitory balance within neural networks. In this process *Ucp2* plays a key role at least in the ventral hippocampus.

Using a test battery of specific behavioral paradigms relevant to hippocampal functions, *Ucp2*^MGKO^ male mice showed heightened anxiety level when comparing to control animals. However, this alteration was not found in female knock-out mice. Interestingly, testing in other behavioral tasks preferentially depending on dCA1 function, such as novel object recognition, marble burying, and Y-maze did not reveal any obvious alteration between *Ucp2*^MGKO^ and control animals of either gender. Overall, these results are in accordance with findings of dysfunctional neural oscillations and plasticity restricted to the vCA1. The hippocampal vCA1 with its direct projections to the prefrontal cortex and further to basolateral amygdala is involved in regulation of various emotional states including anxiety-like behavior (26), (27). Intrinsic deficit in oscillatory activity, as observed in *Ucp2*^MGKO^ male mice, may lead to functional disconnections between these regions causing abnormal behavior outcomes, such as pathological anxiety. In fact, it has been shown that these three regions function as distributed network with high degree of interdependence, and disruption of any element within the network alters anxiety behavior (28), (29), (30), (31).

In this study we observed gender disparity in *Ucp2*^MGKO^ mice with female mice spared from alterations in network function and behavior. Although these differences may be attributed to divergent effects of steroid hormones on neuronal and microglial functions (32), there might be other factors that require detailed analysis in the future.

In summary, we provide evidence of chronotypical and *Ucp2*-mediated crosstalk between microglia and neurons that is involved in the region-specific regulation of hippocampal function and anxiety-like behaviors. This previously unrecognized role of *Ucp2* in maintaining complex behaviors through microglial fitness in the adult brain may open new perspectives for the discovery of novel treatments for psychiatric and neurodegenerative disorders.

## Acknowledgement

This work was supported by NIH grants AG051459, AG052005, AG052986, AG067329 and DK111178 to T.L.H, and, NIH grant DK120321 to S.D. YY was supported by the Cell Science Research Foundation Researcher Training Fellowship, the Uehara Memorial Overseas Research Fellowship (2017-2019), and the Japan Society for the promotion of Science (JSPS) Overseas Research Fellowship (2019–2021).

## Author contribution

Y.Y. and T.L.H. generated the concept for the study. Y.Y., M.S., M.S-P., X-B.G. carried out experiments and analyzed data. J.D.K and S.D. provided the animal model. Y.Y., M.S. and T.L.H. wrote the paper with input from all authors.

## Supplementary material

**Supplemental Fig. 1.**
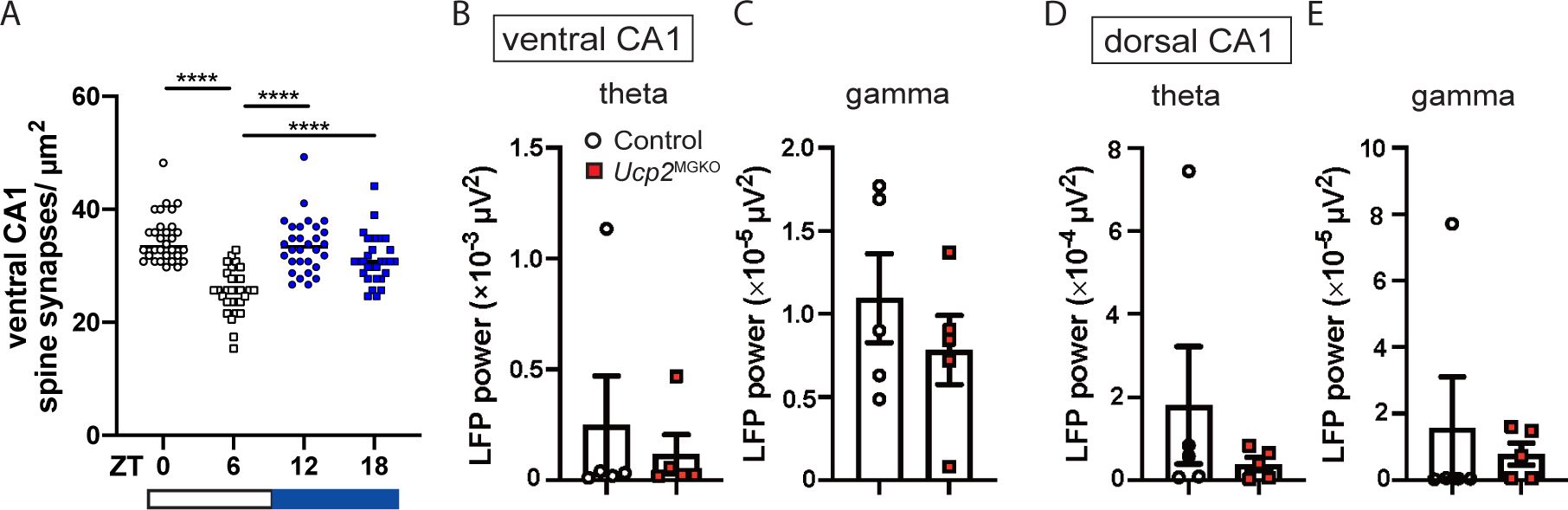
Phasic changes of spine synapse number and hippocampal neurophysiologic characteristics in female mice. **A**, Spine synapse number in female vCA1 at different zeitgeber time periods (ZT0-ZT18). **B-E**, LFPs oscillation power in theta (4-9 Hz) (**B, D**), and gamma (30-90 Hz) (**C, E**) bands from hippocampal vCA1 and dCA1 regions of control and *Ucp2*^*MGKO*^ female mice. Data are presented as scatter plot graph of replicates (n=3 mice/group, 8-12 images per animal) with medians indicated by horizontal lines (**A**), and histograms with individual values and s.e.m. (n=5 mice/group) (**B-E**). ****p<0.0001, by one-way ANOVA with Tukey’s post hoc test.

**Supplemental Fig. 2.**
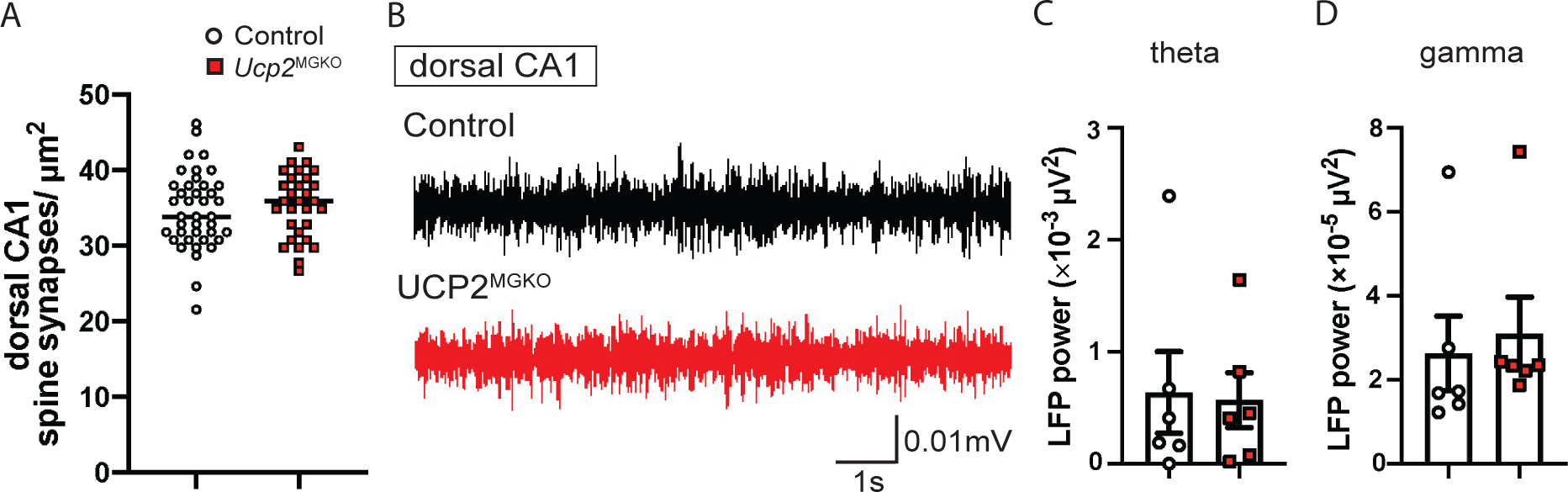
Spine synapse number counts and neurophysiologic characteristics of hippocampal dCA1 in control and *Ucp2*^*MGKO*^ male mice. **A**, Spine synapse number in dCA1 of male control and *Ucp2*^*MGKO*^ mice. **B**, Typical traces of spontaneous local field potentials (LFPs) in gamma frequency range from control (black) and *Ucp2*^*MGKO*^ (red) male mice recorded *in vivo* from dCA1 region. Both signals are digitally band-pass filtered between 30 and 90 Hz. **C-D**, LFPs oscillation power in theta (4-9 Hz) (**C**), and gamma (30-90 Hz) (**D**) bands from dCA1 of control and *Ucp2*^*MGKO*^ male mice. Data are presented as scatter plot graph of replicates (n=3 mice/group, 8-12 images per animal) with medians indicated by horizontal lines (**A**), and histograms with individual values and s.e.m. (n=6 mice/group) (**C, D**).

**Supplemental Fig. 3.**
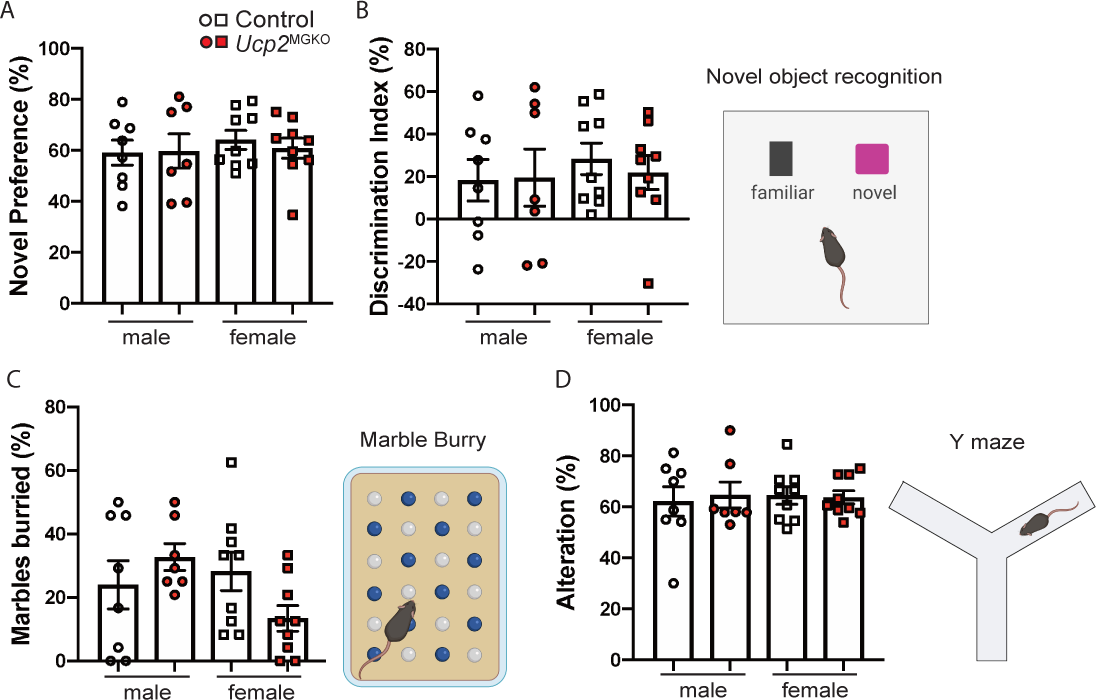
Behavioral tests of control and *Ucp2*^*MGKO*^ mice. **A**,**B**, Novel object recognition test. **C**, Marble-burying test. **D**, Y maze test. Data are presented as histograms with individual values and s.e.m. (n=7-9 mice/group per sex).

